# Heterosynaptic NMDA Receptor Plasticity in Hippocampal Dentate Granule Cells

**DOI:** 10.1101/2022.01.12.476040

**Authors:** Alma Rodenas-Ruano, Kaoutsar Nasrallah, Stefano Lutzu, Maryann Castillo, Pablo E. Castillo

## Abstract

The dentate gyrus is a key relay station that controls information transfer from the entorhinal cortex to the hippocampus proper. This process heavily relies on dendritic integration by dentate granule cells (GCs) of excitatory synaptic inputs from medial and lateral entorhinal cortex via medial and lateral perforant paths (MPP and LPP, respectively). N-methyl-D-aspartate receptors (NMDARs) can contribute significantly to the integrative properties of neurons. While early studies reported that excitatory inputs from entorhinal cortex onto GCs can undergo activity-dependent long-term plasticity of NMDAR-mediated transmission, the input-specificity of this plasticity along the dendritic axis remains unknown. Here, we examined the NMDAR plasticity rules at MPP-GC and LPP-GC synapses using physiologically relevant patterns of stimulation in acute rat hippocampal slices. We found that MPP-GC, but not LPP-GC synapses, expressed homosynaptic NMDAR-LTP. In addition, induction of NMDAR-LTP at MPP-GC synapses heterosynaptically potentiated distal LPP-GC NMDAR plasticity. The same stimulation protocol induced homosynaptic α-amino-3-hydroxy-5-methyl-4-isoxazolepropionic acid receptor (AMPAR)-LTP at MPP-GC but heterosynaptic AMPAR-LTD at distal LPP synapses, demonstrating that NMDAR and AMPAR are governed by different plasticity rules. Remarkably, heterosynaptic but not homosynaptic NMDAR-LTP required Ca^2+^ release from intracellular, ryanodine-dependent Ca^2+^ stores. Lastly, the induction and maintenance of both homo- and heterosynaptic NMDAR-LTP were blocked by GluN2D antagonism, suggesting the recruitment of GluN2D-containing receptors to the synapse. Our findings uncover a mechanism by which distinct inputs to the dentate gyrus may interact functionally and contribute to hippocampal-dependent memory formation.

**Significance Statement:** NMDARs are key players in synaptic plasticity. In addition to their classical role as coincidence detectors and triggers of AMPAR plasticity, there is compelling evidence that NMDARs can undergo activity-dependent plasticity independent of AMPAR plasticity. However, whether NMDAR-plasticity is expressed heterosynaptically remains unclear. Here, in the dentate gyrus of the hippocampus, we show that the induction of burst timing-dependent LTP of NMDAR-mediated transmission at proximal medial perforant path synapses is accompanied by heterosynaptic NMDAR-LTP at lateral perforant path synapses. These findings provide the first evidence for heterosynaptic NMDAR plasticity, which may have important consequences on the dendritic integration of functionally distinct excitatory inputs by dentate granule cells.

## INTRODUCTION

Activity-dependent changes in synaptic strength are widely regarded as a key mechanism underlying experience-driven refinement of neuronal circuits and memory formation (Mayford et al., 2012; Takeuchi et al., 2014). Synaptic plasticity can manifest in multiple forms, including homo- and heterosynaptic plasticity. While homosynaptic plasticity is expressed at active synapses only, heterosynaptic plasticity involves a change in strength at neighboring naive synapses. Although it is well-established that glutamate NMDA receptors (NMDARs) mediate excitatory synaptic transmission and can themselves undergo long-term potentiation (LTP) and long-term depression (LTD) of NMDAR-mediated transmission (Rebola et al., 2010; Hunt and Castillo, 2012), most studies have focused on homosynaptic (Herring and Nicoll, 2016; Diering and Huganir, 2018) and heterosynaptic (Chistiakova et al., 2014; Chater and Goda, 2021) plasticity of the AMPA receptor (AMPAR)-mediated component of glutamatergic excitatory transmission. However, whether NMDAR-mediated transmission undergoes heterosynaptic plasticity has not yet been investigated.

The dentate gyrus is the main input area of the hippocampus (Amaral et al., 2007; Jonas and Lisman, 2014). Dentate gyrus granule cells (GCs) receive prominent cortical projections from the medial and lateral entorhinal cortex which convey context and content related information, through the medial- and lateral perforant pathway (MPP and LPP), respectively (Knierim et al., 2014). Activity-dependent changes of these inputs, and their dendritic integration by GCs are critical to dentate gyrus information processing (Schmidt-Hieber et al., 2007; Krause et al., 2008; Krueppel et al., 2011; Knierim et al., 2014). MPP and LPP inputs onto GCs express robust homosynaptic (Bliss and Lomo, 1973; Colino and Malenka, 1993; Richter-Levin et al., 1995) and heterosynaptic (Abraham et al., 2007; Jungenitz et al., 2018) AMPAR long-term plasticity. Furthermore, early studies showed that high frequency stimulation can also trigger homosynaptic NMDAR plasticity at MPP-GC excitatory synapses (O’Connor et al., 1994; Harney et al., 2006). To our knowledge, whether homo- or heterosynaptic NMDAR plasticity is expressed at distal LPP inputs onto GCs is unknown.

Here, we examined homo- and heterosynaptic plasticity of NMDAR-mediated transmission at entorhinal-GC synapses by pairing physiologically relevant patterns of pre-and postsynaptic burst activity in acute rat and mouse hippocampal slices. Burst pairing stimulation induced homosynaptic NMDAR-LTP at MPP-GC but not at LPP-GC synapses. In addition, homosynaptic NMDAR-LTP at MPP-GCs synapses was accompanied by heterosynaptic NMDAR-LTP at non-stimulated distal LPP-GC synapses. Conversely, homosynaptic AMPAR-LTP at MPP-GC synapses was accompanied by heterosynaptic AMPAR-LTD at LPP-GC synapses. Both homo- and heterosynaptic NMDAR-LTP were blocked by GluN2D antagonism. Thus, our study provides the first direct evidence of heterosynaptic LTP of NMDAR-mediated transmission. This novel form of NMDAR-LTP may contribute to dentate gyrus-dependent forms of memory.

## METHODS

### Experimental Model and Subject Details

Postnatal day 19 (P19) to P29 Sprague-Dawley rats and C57BL/6 mice P30 to P60 of either sex were used for electrophysiological experiments. All animals were group housed in a standard 12 hr light/12 hr dark cycle. Animal handling and use followed a protocol approved by the Animal Care and Use Committee of Albert Einstein College of Medicine, in accordance with the National Institutes of Health guidelines.

### Hippocampal slice preparation

Acute transverse hippocampal slices (300-400 μm thick) were prepared from Sprague-Dawley rats and C57BL/6 mice (300 μm thick). Briefly, the hippocampi were isolated and cut using a VT1200s microsclicer (Leica Microsystems Co.) in a solution containing (in mM): 215 sucrose, 2.5 KCl, 26 NaHCO_3_, 1.6 NaH_2_PO_4_, 1 CaCl_2_, 4 MgCl_2_, 4 MgSO_4_ and 20 D-glucose. At 30 min post sectioning, the cutting medium was gradually switched to extracellular artificial cerebrospinal (ACSF) recording solution containing (in mM): 124 NaCl, 2.5 KCl, 26 NaHCO_3_, 1 NaH_2_PO_4_, 2.5 CaCl_2_, 1.3 MgSO_4_ and 10 D-glucose. Slices were incubated for at least 45 min at room temperature in the ACSF solution before recording.

### Electrophysiology

All experiments, unless otherwise stated, were performed at 28 ± 1°C in a submersion-type recording chamber perfused at ∼2 mL min^−1^ with ACSF. Whole-cell patch-clamp recordings using a Multiclamp 700A amplifier (Molecular Devices) were made from GCs voltage clamped at −45 mV (unless otherwise stated) using patch-type pipette electrodes (∼3-4 MΩ) containing (in mM): 135 K-Gluconate, 5 KCl, 0.1 EGTA, 0.04 CaCl_2_, 5 NaOH, 5 NaCl, 10 HEPES, 5 MgATP, 0.4 Na_3_GTP, and 10 D-glucose, pH 7.2 (280-290 mOsm). Series resistance (∼7-25 MΩ) was monitored throughout all experiments with a −5 mV, 80 ms voltage step, and cells that exhibited a significant change in series resistance (> 20%) were excluded from analysis.

In order to stimulate MPP and LPP inputs to GCs, stimulating patch-type pipettes were placed in the middle third of the medial molecular layer, and in the distal part of the outer molecular layer of the dentate gyrus, respectively. To elicit synaptic responses, monopolar square-wave voltage or current pulses (100–200 μs pulse width, 4-25 V or 20-100 μA) were delivered through a stimulus isolator (Isoflex, AMPI, or Digitimer DS2A-MKII) connected to a broken tip (∼10–20 μm) stimulating patch-type micropipette filled with ACSF. Typically, stimulation was adjusted to obtain comparable magnitude synaptic responses across experiments; e.g., 40-100 pA NMDAR-EPSCs (Vh −45 mV). Burst-timing NMDAR plasticity was typically induced in current clamp mode (Vrest ∼ -60 to -70 mV) by pairing presynaptic bursts (6 pulses at 50 Hz) and postsynaptic burst of action potentials (5 action potentials at 100 Hz) delivered at 10 ms interval, for 100 times at 2 Hz.

### Data analysis

Electrophysiological data were acquired at 5 kHz, filtered at 2.4 kHz, and analyzed using custom-made software for IgorPro (Wavemetrics Inc.). The magnitude of LTP was determined by comparing 10 min baseline responses with responses 20-30 min after LTP induction.

### Reagents

Reagents were bath applied following dilution into ACSF from stock solutions stored at -20 °C prepared in water or DMSO, depending on the manufacturers’ recommendation. Final DMSO concentration was < 0.01% total volume. Cyclopiazonic acid, heparin, ryanodine, DQP-1105, QNZ46, D-APV, and MPEP were purchased from Tocris Biosciences. NBQX was purchased from Cayman Chemical Co, while BAPTA, picrotoxin and all salts for making ACSF and internal solutions were purchased from Sigma-Millipore.

### Quantification and Statistical Analysis

Statistical analysis was performed using OriginPro software (OriginLab). The normality of distributions was assessed using the Shapiro-Wilk test. In normal distributions, Student’s unpaired and paired two-tailed t tests were used to assess between-group and within-group differences, respectively. The non-parametric paired sample Wilcoxon signed rank test and Mann-Whitney’s U test were used in non-normal distributions. Statistical significance was set to p < 0.05 (*** indicates p < 0.001, ** indicates p < 0.01, and * indicates p < 0.05). All values are reported as the mean ± SEM.

## RESULTS

### MPP-GC synapses express homosynaptic burst timing-dependent NMDAR-LTP

Burst timing-dependent plasticity (BTDP) of NMDAR-mediated transmission has been reported in the midbrain (Harnett et al., 2009) and in area CA3 of the hippocampus (Hunt et al., 2013), whereas early work utilized high frequency stimulation of MPP axons to elicit NMDAR-LTP at MPP-GC synapses (O’Connor et al., 1994). BTDP mimics *in vivo* activity in the entorhinal cortex (Latuske et al., 2015; Ebbesen et al., 2016; Csordas et al., 2020) and dentate gyrus (Pernia-Andrade and Jonas, 2014; Diamantaki et al., 2016; Senzai and Buzsaki, 2017), so we aimed to test whether pre (MPP axons) – post (GC) burst pairing stimulation could trigger homosynaptic LTP of NMDAR-mediated transmission at MPP-GC synapses. To isolate NMDAR-mediated transmission, whole-cell patch-clamp recordings of GCs (holding potential [Vh] = -45 mV) were performed in the presence of 10 µM NBQX and 100 µM picrotoxin to block AMPAR- and GABA_A_ receptor-mediated transmission, respectively. NMDAR excitatory postsynaptic currents (EPSCs) were evoked using electrical stimulation with a micropipette placed in the medial molecular layer (MML) of the dentate gyrus (see Methods). After 10 minutes of baseline response, a presynaptic (6 pre pulses at 50 Hz) – postsynaptic (5 post pulses at 100 Hz) burst firing pairing protocol with a 10 ms interval, repeated 100 times, was delivered in current clamp mode (holding potential [Vh] = ∼-60 to -70 mV). We found that pre-post burst stimulation induced a robust homosynaptic NMDAR-LTP at MPP-GC synapses, whereas a post-pre pairing sequence had no long-term effect on NMDAR-mediated synaptic transmission (Fig. 1A). In addition, presynaptic and postsynaptic bursts alone did not elicit any NMDAR plasticity (Fig. 1B). Altogether, these results revealed that physiologically relevant patterns of activity trigger robust homosynaptic NMDAR-LTP at MPP-GC synapses, and that burst timing-induced LTP is restricted to pre-post pairings.

**Figure 1:**
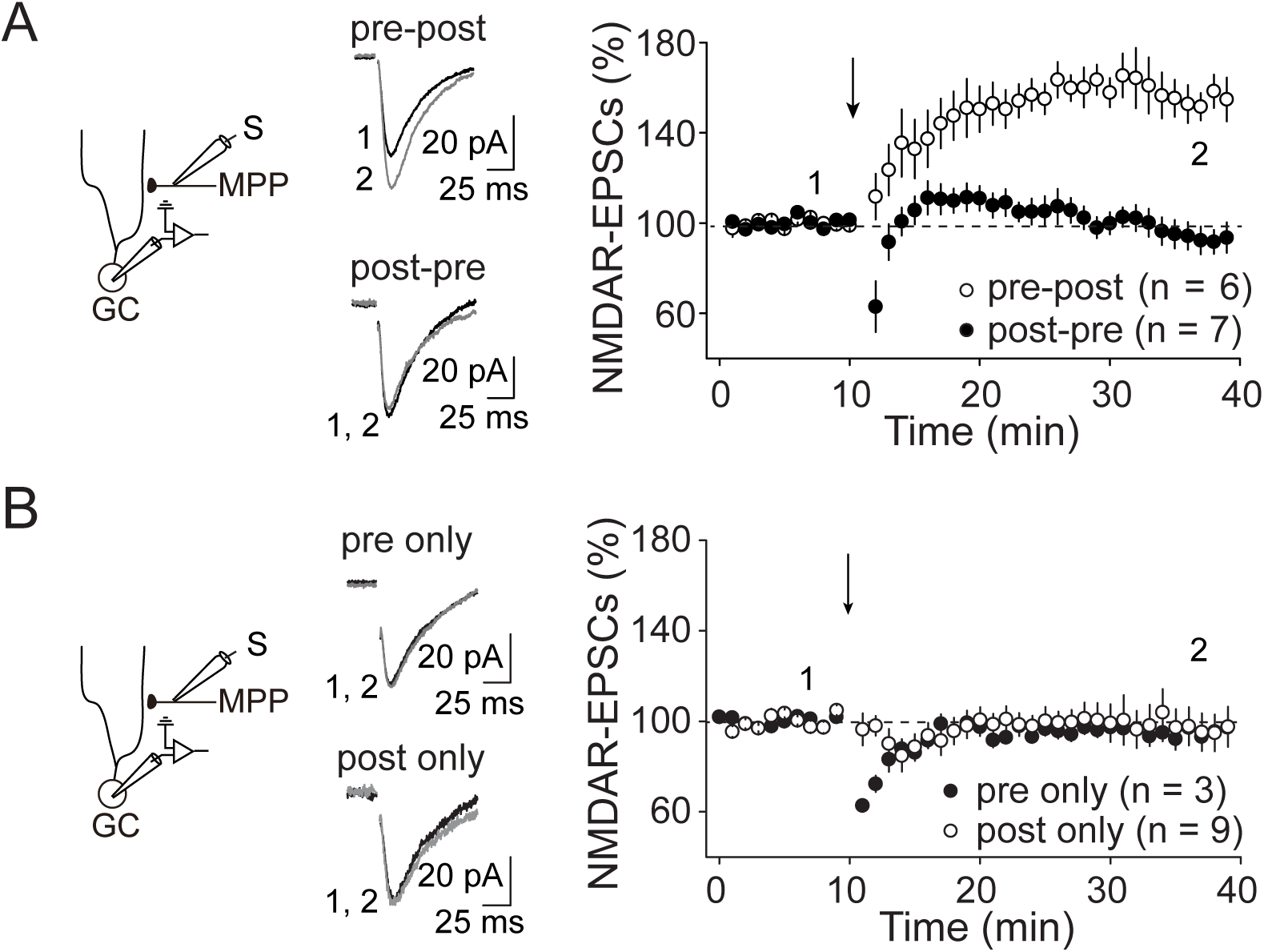
MPP-GC synapses express homosynaptic burst timing-dependent NMDAR-LTP. (A) Left, Diagram illustrating the recording configuration. MPP NMDAR EPSCs were recorded from GC and evoked with a stimulating electrode placed in the middle molecular layer (MML) of the dentate gyrus. Middle, Representative average traces before (1) and after (2) pairing protocol application. Right, Time-course summary plot showing how pairing pre- and postsynaptic activity (pre-post: 6 pre pulses at 50 Hz followed by 5 post pulses at 100 Hz with a 10 ms interval, repeated 100 times every 0.5 s) induced a robust NMDAR-LTP at MPP-GC synapses (white circles, 159.5 ± 7.3 %, p < 0.001, n = 6, paired t-test). In contrast, post-pre protocol (post-pre: 5 post pulses at 100 Hz followed by 6 pre pulses at 50 Hz with a 10 ms interval, repeated 100 times every 0.5 s) resulted did not change NMDAR EPSC amplitude (black circles, 96.9 ± 5.3 %, p = 0.58, n = 7, paired t-test). Pairing protocol was delivered at the time point indicated by the vertical arrow. (B) Representative traces and summary plot showing how neither presynaptic (black circles, pre only, p = 0.51, n = 3, paired t-test) nor postsynaptic (white circles, post only, p = 0.89, n = 9, paired t-test) bursts alone elicited any long-lasting change in NMDAR EPSC amplitude. Numbers in parentheses indicate the number of cells. Summary data represent mean ± s.e.m.

Previous work demonstrated that NMDAR-LTP at CNS synapses (Hunt and Castillo, 2012), including MPP-GC synapses (O’Connor et al., 1994; Harney et al., 2006), relies on a postsynaptic mechanism of induction that includes NMDAR and type 5 metabotropic glutamate receptor (mGluR5) co-activation and a rise in postsynaptic [Ca^2+^]. We found that bath application of the mGluR5 antagonist MPEP (4 µM) (Fig. 2A) and intracellular loading of the Ca^2+^ chelating agent BAPTA (10 mM) abolished LTP (Fig. 2B), indicating that both mGluR5 and postsynaptic calcium are also required for pre-post burst-induced NMDAR LTP at MPP-GC synapses.

**Figure 2:**
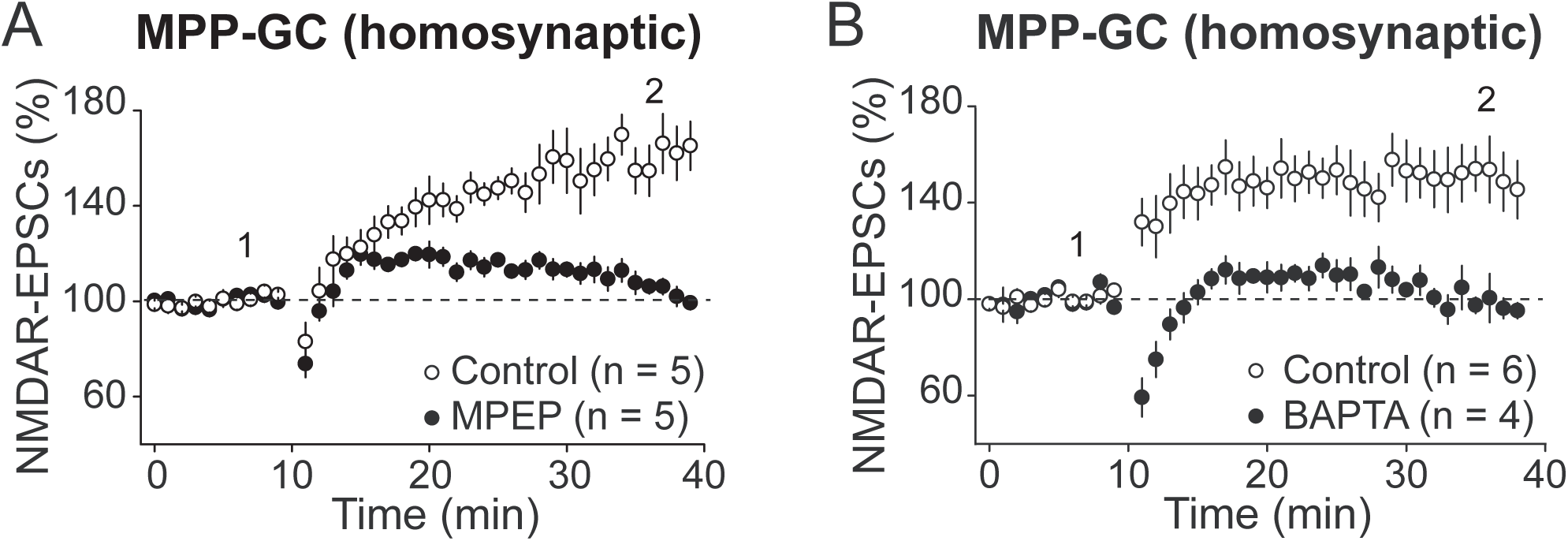
MPP-GC burst timing-dependent NMDAR-LTP requires mGluR5 activation and postsynaptic calcium. (A) Bath application of the mGluR5 antagonist MPEP (4 µM) significantly reduced homosynaptic NMDAR LTP (black circles, 107.6 ± 2.4 %, n = 5) as compared to interleaved controls (white circles, 154.3 ± 9.5 %, p < 0.01, n = 5, paired t-test; MPEP vs control: p < 0.01, unpaired t-test). (B) Intracellular loading of the Ca^2+^ chelating agent BAPTA (10 mM) abolished NMDAR-LTP (black circles, 98.9 ± 3.4 %, p = 0.76, n = 4, paired t-test) relative to controls (white circles, 151.7 ± 11.4 %, p < 0.01, n = 6, paired t-test; BAPTA vs control: p < 0.01, unpaired t-test). Numbers in parentheses indicate the number of cells. Summary data represent mean ± s.e.m.

### LPP-GC synapses express robust heterosynaptic NMDAR LTP but heterosynaptic AMPAR LTD

We next tested whether induction of burst timing-dependent NMDAR-LTP at MPP-GC synapses could heterosynaptically induce plasticity at distal LPP-GC synapses. Some forms of heterosynaptic plasticity are compensatory in nature –e.g., homosynaptic LTP can result in heterosynaptic LTD at neighboring synapses (Chistiakova et al., 2014; Chater and Goda, 2021). To test for NMDAR heterosynaptic plasticity, NMDAR-EPSCs were monitored in the same GC in response to MPP and LPP stimulation, by placing stimulation micropipettes in MML and the most distal border of the outer molecular layer (OML) of the dentate gyrus, respectively. Both stimulation pipettes were placed on the same side of the GC to activate similar dendritic branches. After a 10-min baseline recording, pre-post burst stimulation was applied at MPP-GC synapses as previously described (Figs. 1 and 2). Surprisingly, we found that homosynaptic MPP-GC NMDAR-LTP was accompanied by heterosynaptic NMDAR-LTP at LPP-GC synapses (Fig. 3A and B). Because single dendritic branches perform local computations and may act as fundamental functional units of a neuron (Branco and Hausser, 2010), we hypothesized that the postsynaptic signaling that mediates heterosynaptic plasticity could be restricted to the dendritic branches expressing homosynaptic LTP. To test this possibility, we placed the LPP stimulation pipette on the other side of the recorded GC, which is expected to activate LPP inputs onto different dendritic branches. We found that these LPP inputs did not express any heterosynaptic plasticity (Fig. 3C). Moreover, application of the pre-post burst stimulation protocol at LPP-GC synapses did not induce any form of homosynaptic NMDAR plasticity (Fig. 3D). To test whether homosynaptic MPP-GC NMDAR-LTP and heterosynaptic LPP-GC NMDAR-LTP could also be observed in mice, we repeated the experiment reported in figure 3A,B in acute mouse hippocampal slices. Both forms of NMDAR plasticity were also present in mice (Fig. 3E), suggesting a conserved plasticity mechanism. Altogether, these results revealed that MPP and LPP synaptic inputs onto GCs undergo distinct forms of NMDAR plasticity. While pre-post burst stimulation induced homosynaptic NMDAR-LTP at MPP inputs, only heterosynaptic NMDAR-LTP was observed at LPP inputs.

**Figure 3:**
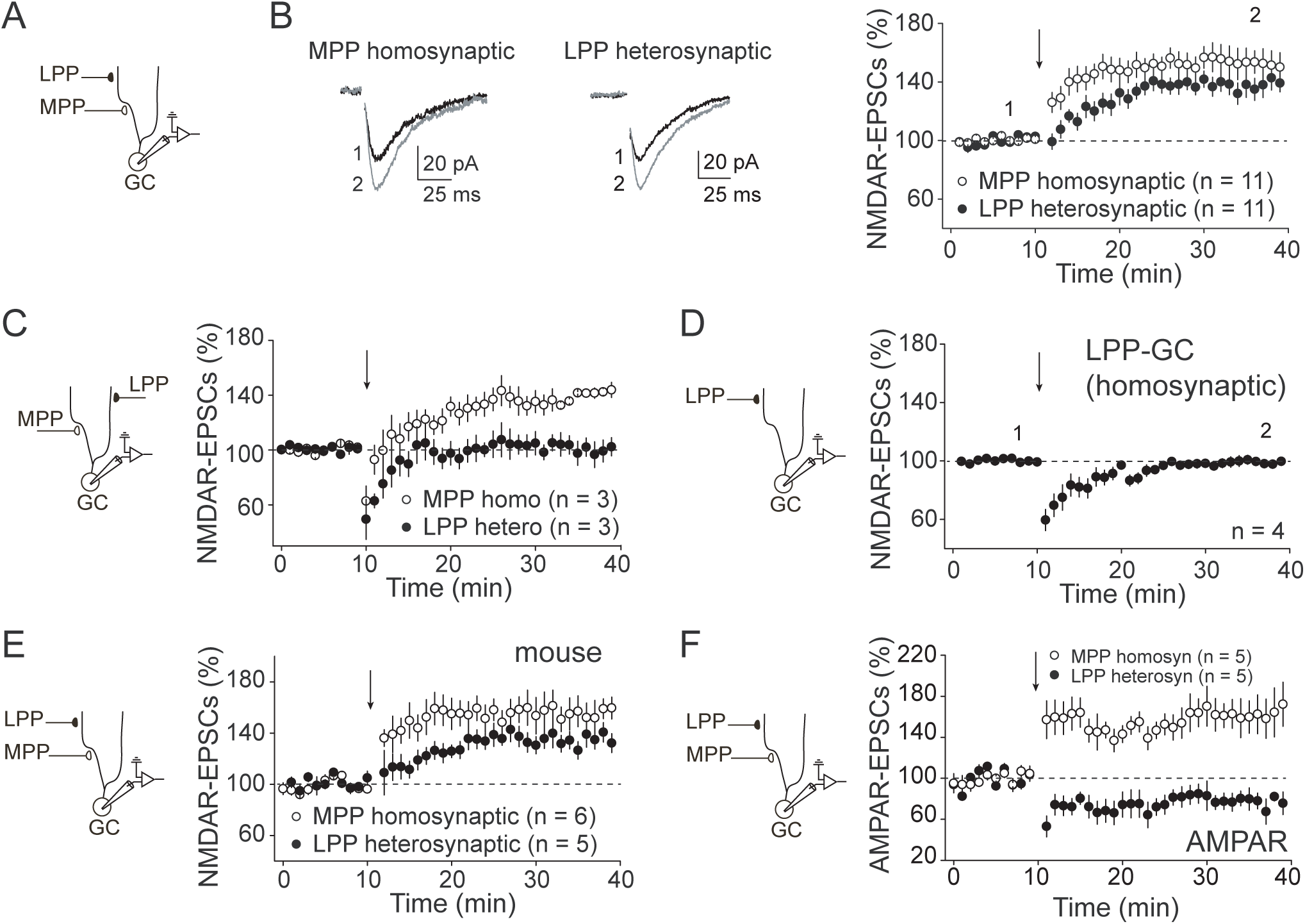
LPP-GC synapses express heterosynaptic NMDAR-LTP but heterosynaptic AMPAR-LTD. (A) Left, Diagram illustrating the recording configuration. MPP and LPP NMDAR EPSCs were recorded from the same GC. The LPP stimulation micropipette was placed in the most distal border of the outer molecular layer, directly above the MPP stimulation micropipette, which was positioned within the MML of the dentate gyrus. (B) Left, Representative average traces. Right, Time-course summary plot showing how pre-post pairing protocol induced at MPP (6 pre pulses at 50 Hz followed by 5 post pulses at 100 Hz with a 10 ms interval, repeated 100 times every 0.5 s), resulted in homosynaptic NMDAR-LTP at MPP-GC synapses (white circles, 153.1 ± 7.9 %, p < 0.001, n = 11, paired t-test), accompanied by heterosynaptic NMDAR-LTP at distal LPP-GC synapses (black circles, 137.2 ± 5.1 %, p < 0.001, n = 11, paired t-test). (C) Left, diagram illustrating recording configuration. MPP and LPP NMDAR EPSCs were recorded from the same GC. LPP stimulation micropipette was placed on the distal OML, at least 250 µm lateral from the location of the MPP stimulation micropipette. Right, summary plot showing that pre-post pairing elicited homosynaptic LTP at MPP-GC synapses (white circles, 138.4 ± 3.7 %, p < 0.01, n = 3, paired t-test) while it did not trigger any change in heterosynaptic LPP NMDAR EPSC amplitude under these conditions (black circles, 101.4 ± 6.6 %, p = 0.85, n = 3, paired t-test).). (D) Left, Diagram illustrating the recording configuration. LPP NMDAR EPSCs were recorded from GC and evoked with stimulating electrodes placed in OML of the dentate gyrus. Right, summary data showing how Pre-post pairing protocol failed to elicit homosynaptic NMDAR LTP at LPP-GC synapses (98.1 ± 4.6 %, p = 0.71, n = 4, paired t-test). (E) Summary data obtained from experiments performed in acute mouse hippocampal slices, shows that heterosynaptic NMDAR LTP at DGCs is a conserved phenomenon. Pre-post pairing protocol induced at MML, resulted in homosynaptic NMDAR-LTP at MPP-GC synapses (white circles, 155.8 ± 11.3 %, p < 0.01, n = 6, paired t-test), accompanied by heterosynaptic LTP at distal LPP-GC synapses (black circles, 134.5 ± 6.8 %, p < 0.01, n = 5, paired t-test). (F) Left, Diagram illustrating the recording configuration. MPP and LPP AMPAR EPSCs were recorded from the same GC. Right, Time-course summary plot shows that pre-post pairing protocol at MPP inputs induced homosynaptic AMPAR-LTP at MPP-GC synapses (white circles, 163.2 ± 15.8 %, p < 0.05, n = 5, paired t-test), while it triggered heterosynaptic AMPAR-LTD at LPP-GC synapses (black circles, 77.7 ± 7.6 %, p < 0.05, n = 5, paired t-test). Numbers in parentheses indicate the number of cells. Summary data represent mean ± s.e.m.

We wondered whether the homosynaptic and heterosynaptic BTDP we observed was unique to NMDARs, or it could also be expressed by AMPARs. While virtually nothing was known about NMDAR heterosynaptic plasticity, several reports showed that AMPARs can undergo heterosynaptic changes in neighboring synapses (Chistiakova et al., 2014; Chater and Goda, 2021). Yet, only a handful of *in vivo* studies with mixed results have investigated heterosynaptic AMPAR plasticity in the dentate gyrus (Abraham et al., 2007; Bromer et al., 2018; Jungenitz et al., 2018) and, to our knowledge, no *in vitro* study has characterized any form of functional heterosynaptic AMPAR plasticity at LPP-GC synapses. To address this knowledge gap, we monitored AMPAR-EPSCs at both MPP- and LPP-GC synapses in the same GC (Vh = -60 mV), in presence of the GABA_A_ receptor antagonist picrotoxin (100 uM). After establishing a baseline, we applied pre-post burst stimulation at MPP-GC synapses and found that AMPAR-EPSCs express homosynaptic LTP at MPP-GC synapses but heterosynaptic AMPAR-LTD at LPP-GC synapses (Fig. 3F). These results suggest that BTDP modifies the relative contribution of AMPA and NMDA receptors at MPP and LPP synaptic inputs.

### Internal Ca^2+^ stores are required for heterosynaptic but not homosynaptic NMDAR-LTP

Previous work by our lab showed that internal Ca^2+^ stores contribute to the expression of homosynaptic NMDAR-LTP at mossy fiber to CA3 pyramidal cell synapses (Hunt et al., 2013). We then tested the role of internal Ca^2+^ stores in both homosynaptic and heterosynaptic NMDAR-LTP at GC synapses. To this end, we used the cell-permeable inhibitor of sarcoplasmic reticulum Ca^2+^-ATPase, cyclopiazonic acid (CPA 30 µM, 40-50 min pre-incubation, also included in the perfusion) to deplete internal Ca^2+^ stores. We found that CPA abolished the induction of LPP-GC heterosynaptic plasticity (Fig. 4A) but had no effect on MPP-GC homosynaptic plasticity (Fig. 4B). Internal Ca^2+^ can be released from endoplasmic reticulum (ER) stores via IP3 signaling or ryanodine receptor activation. To determine which of these mechanisms is required for heterosynaptic NMDAR-LTP, we loaded GCs with heparin (2 mg/ml) to block IP3-mediated Ca^2+^ release, or incubated (40-50 min) and perfused hippocampal slices with ryanodine (100 µM) to block ryanodine receptor-mediated Ca^2+^ release. We found that induction of heterosynaptic LPP-GC NMDAR-LTP was abolished by ryanodine (Fig. 4C) but normally induced in presence of heparin (fig. 4C), suggesting that heterosynaptic LTP requires ryanodine but not IP3 receptors. In contrast, homosynaptic NMDAR-LTP at MPP-GC was normally induced in presence of both ryanodine and heparin (Fig. 4D), indicating that the rise in postsynaptic Ca^2+^ at MPP-GC synapses is independent of internal Ca^2+^ stores. Taken together, these results show that homosynaptic vs. heterosynaptic NMDAR-LTP have unique Ca^2+^ requirements for induction, where internal Ca^2+^ release is required only for heterosynaptic NMDAR-LTP at LPP-GC synapses. By triggering ryanodine receptor-dependent release of Ca^2+^ from internal stores, homosynaptic MPP-GC NMDAR-LTP could change LPP-GC synaptic plasticity rules.

**Figure 4.**
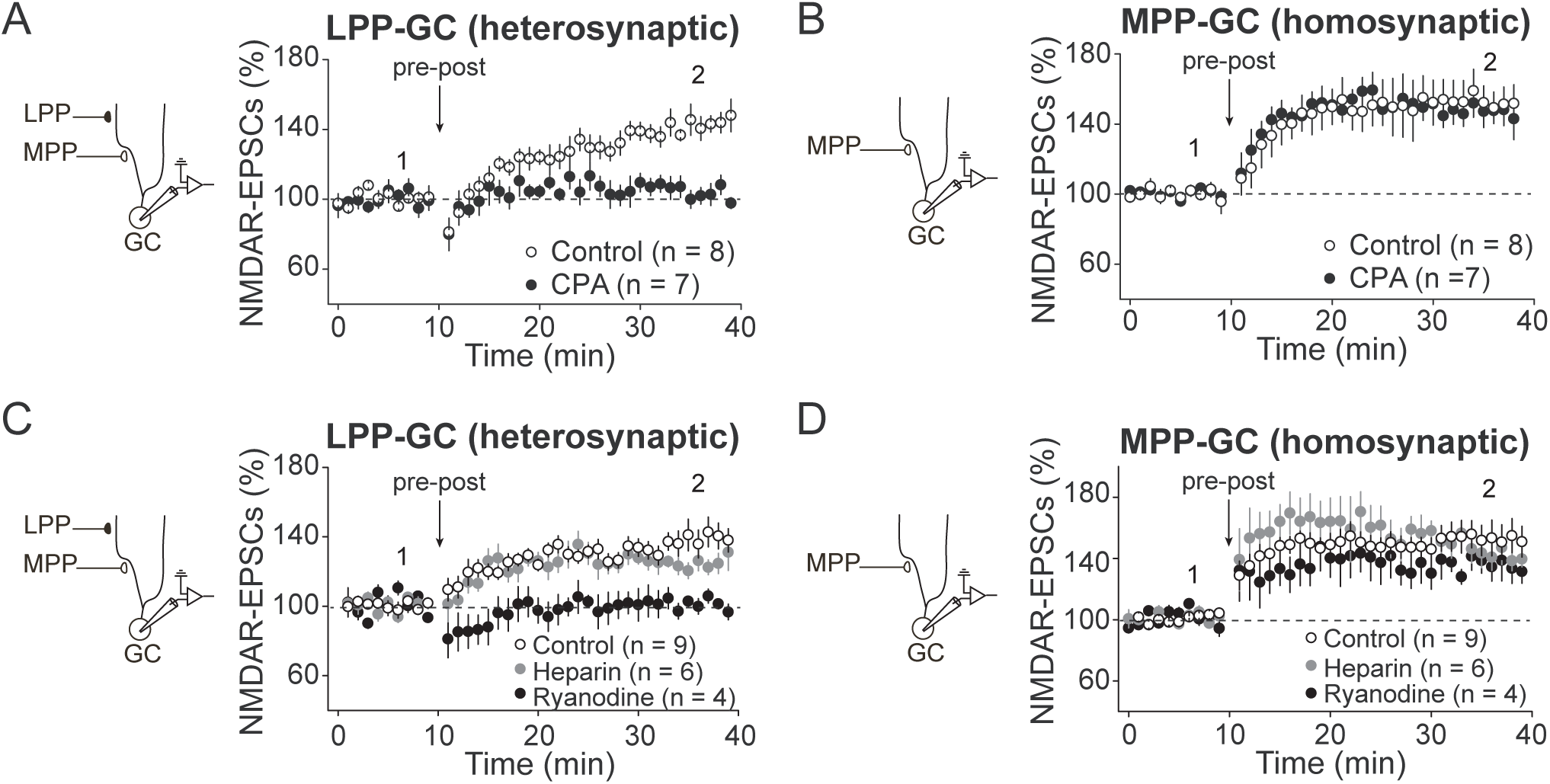
Internal calcium stores are required for heterosynaptic but not homosynaptic NMDAR-LTP. (A) Left, Diagram illustrating the recording configuration. LPP NMDAR EPSCs were recorded from GC and pre-post pairing protocol was delivered at MPP inputs. Application of the sarcoplasmic reticulum Ca^2+^-ATPase inhibitor CPA (30 µM, 40-50 min pre-incubation and bath application) completely abolished heterosynaptic NMDAR-LTP at LPP-GC synapses (black circles, 106.6 ± 3.8 %, p = 0.13, n = 7, paired t-test), relative to interleaved controls (white circles, 140.5 ± 4.2 %, p < 0.001, n = 8, paired t-test, CPA vs control: p < 0.001, unpaired t-test). (B) MPP NMDAR EPSCs were recorded from GC and pre-post pairing protocol was delivered at MPP inputs. CPA application (30 µM, 40-50 min pre-incubation and bath application) had no effect on homosynaptic NMDAR-LTP at MPP-GC synapses (black circles, 149.4 ± 6.5 %, p < 0.001, n = 7, paired t-test, CPA vs control: p < 0.001, unpaired t-test) as compared to interleaved controls (white circles, 151.3 ± 7.8 %, p < 0.001, n = 8, paired t-test, CPA vs control: p = 0.75, unpaired t-test). (C) LPP NMDAR EPSCs were recorded from GC and pre-post pairing protocol was delivered at MPP inputs. GCs were loaded with heparin (2 mg/ml) to block IP3-mediated Ca^2+^ release, or slices were pre-treated (40-50 min prior to induction) and perfused during experiment with ryanodine (100 µM) to block ryanodine receptor-mediated Ca^2+^ release. Induction of heterosynaptic plasticity was completely blocked by ryanodine (black circles, 101.4 ± 4.1 %, p = 0.76, n = 4, paired t-test, ryanodine vs control: p < 0.05, unpaired t-test), but was comparable to controls (white circles, 135.4 ± 5.6 %, p < 0.001, n = 9, paired t-test) in the presence of heparin (gray circles, 124.2 ± 4.0 %, p < 0.01, n = 6, paired t-test, heparin vs control: p = 0.44, unpaired t-test). (D) MPP NMDAR EPSCs were recorded from GC and pre-post pairing protocol was delivered at MPP inputs. Inhibition of ryanodine (black circles, 135.3 ± 8.7 %, p < 0.05, n = 4, paired t-test; ryanodine vs control: p = 0.46, unpaired t-test) or IP3 (gray circles, 146.5 ± 7.1 %, p < 0.01, n = 6, paired t-test; heparin vs control: p = 0.44, unpaired t-test) receptors had no effect on homosynaptic NMDAR-LTP at MPP-GC synapses compared to interleaved controls (white circles, 152.6 ± 8.9 %, p < 0.001, n = 9, paired t-test). Numbers in parentheses indicate the number of cells. Summary data represent mean ± s.e.m.

### Homosynaptic and heterosynaptic NMDAR-LTP are likely mediated by the synaptic recruitment of GluN2D-containing receptors

Previous studies reported that HFS-induced homosynaptic NMDAR-LTP at MPP-GC synapses is mediated by the recruitment of GluN2D-containing NMDARs to the synapse (Lozovaya et al., 2004; Harney et al., 2008). To assess the role of GluN2D subunits in burst timing-dependent homosynaptic and heterosynaptic NMDAR-LTP, we used two different non-competitive, activity-dependent GluN2C/GluN2D selective antagonists, DQP-1105 and QNZ46 (Hansen and Traynelis, 2011; Monaghan et al., 2012). Although these antagonists may not distinguish between GluN2C- and GluN2D-containing NMDARs, GluN2C expression in the adult hippocampus (Sanz-Clemente et al., 2013), and neurons in general (Alsaad et al., 2019), is negligible. We first tested the effect of DQP-1105 on both homosynaptic MPP-GC and heterosynaptic LPP-GC NMDAR-LTP, and found that bath application of 30 µM DQP-1105 abolished both homosynaptic and heterosynaptic plasticity (Fig. 5A,B). Most evidence indicates that GluN2D-containing receptors are mainly extrasynaptic in adult animals (Cull-Candy et al., 2001; Paoletti et al., 2013). Consistent with this notion, bath application of the antagonists DQP-1105 or QNZ46 (30 µM) had no effect on MPP- and LPP-GC basal synaptic transmission, whereas the non-selective NMDAR antagonist D-APV abolished these responses (50 µM) (Fig. 5C,D). Using not so selective pharmacology for GluN2D-containing receptors, it has been suggested that these receptors could be recruited to the synapse upon activity and mediate NMDAR-LTP (Harney et al., 2008). If so, the contribution of GluN2D-containing receptors to NMDAR-mediated transmission should increase after LTP induction. Remarkably, application of the selective GluN2D antagonists (DQP-1105 or QNZ46, 30 µM) 30 min after the induction of MPP-GC NMDAR-LTP significantly decreased both MPP-GC and LPP-GC NMDAR-mediated transmission (Fig. 5E,F). These data strongly suggest that both homo and heterosynaptic NMDAR-LTP in the dentate gyrus are due to the recruitment of GluN2D-containing receptors to the synapse.

**Figure 5.**
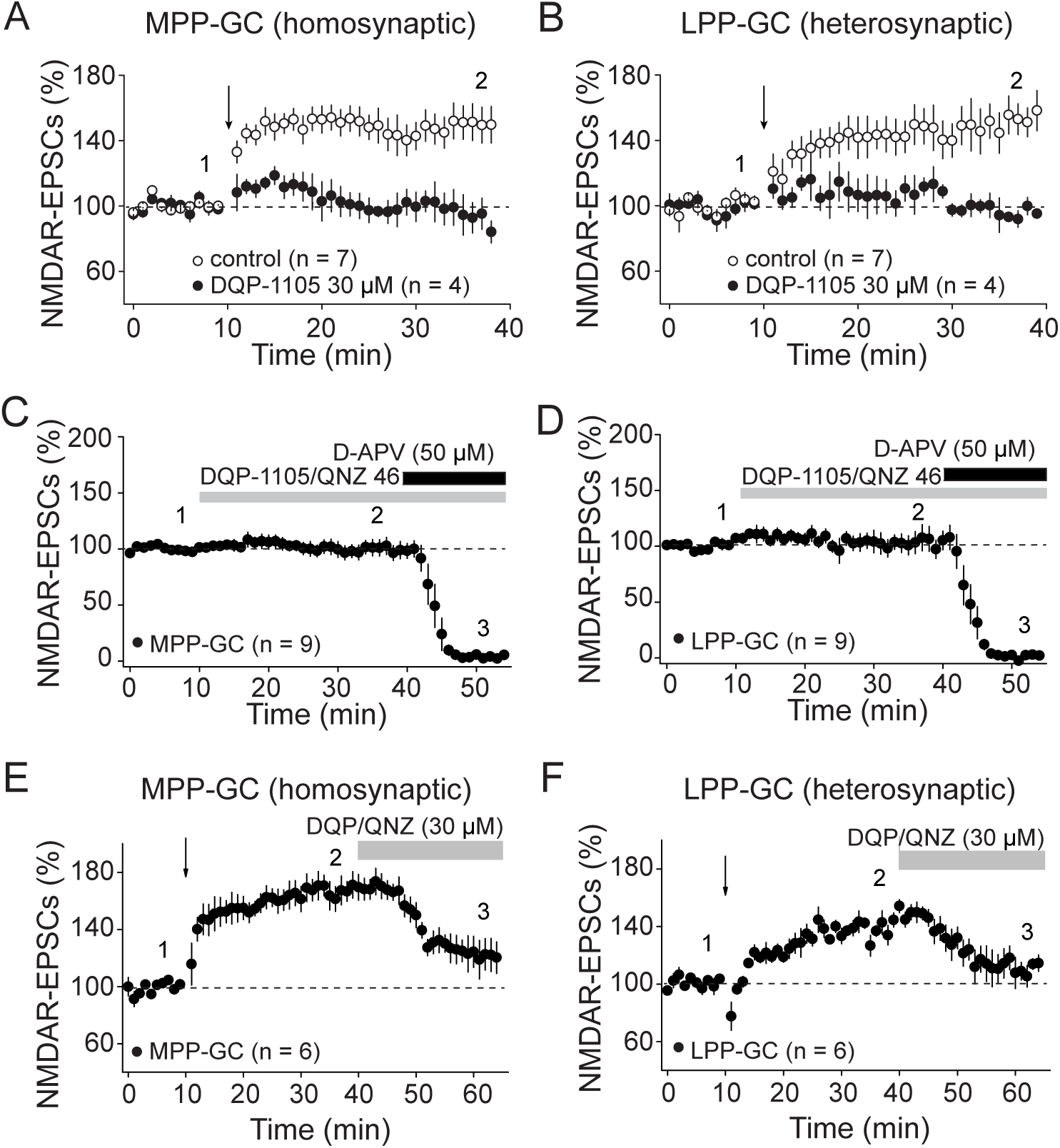
Homosynaptic and heterosynaptic NMDAR-LTP are likely due to GluN2D recruitment to the synapse. (A) Bath application of the GluN2D antagonist DQP-1105 (30 µM) completely abolished homosynaptic NMDAR-LTP at MPP-GC synapses (control: 149.1 ± 8.1 %, p < 0.001, n = 7, paired t-test; DQP-1105: 95.6 ± 8.8 %, p = 0.65, n = 4, paired t-test, DQP-1105 vs control: p < 0.01, unpaired t-test). (B) Induction of heterosynaptic NMDAR-LTP at LPP-GC synapses was also blocked in the presence of 30 µM DQP-1105 (control: 147.8 ± 9.1 %, p < 0.01, n = 7, paired t-test; DQP-1105: 99.1 ± 2.2 %, p = 0.71, n = 4, paired t-test, DQP-1105 vs control: p < 0.01, unpaired t-test). (C, D) Bath application of DQP-1105 or QNZ46 had no effect on basal NMDAR synaptic transmission at either MPP-GC (C, 100.2 ± 5.9 %, p = 0.98, n = 9, paired t-test) or LPP-GC synapses (D, 103.4 ± 7.7 %, p = 0.67, n = 9, paired t-test). Application of D-APV abolished these EPSCs, confirming these responses were mediated by NMDARs (C, MPP: 5.3 ± 1.6 %, p < 0.0001, n = 5, paired t-test; D, LPP: 9.0 ± 3.7 %, p < 0.001). (E, F) Pre-post pairing LTP induction protocol was delivered at MPP inputs. Bath application of either DQP-1105 (30 µM) or QNZ46 (30 µM) 30 min after NMDAR-LTP induction substantially reduced both homosynaptic (E, DQP-1105 vs post-LTP: p < 0.05, unpaired t-test) and heterosynaptic (F, DQP-1105 vs post-LTP: p < 0.05, unpaired t-test) NMDAR-EPSC amplitude. Numbers in parentheses indicate the number of cells. Summary data represent mean ± s.e.m.

## DISCUSSION

In this study, we discovered that pairing physiologically relevant patterns of pre and postsynaptic burst activity induced homosynaptic LTP of NMDAR-mediated transmission at MPP-GC synapses. Unexpectedly, this BTDP also triggered robust heterosynaptic NMDAR-LTP at non-stimulated LPP-GC synapses. In contrast, the same induction protocol that triggered homo- and heterosynaptic NMDAR-LTP, elicited homosynaptic MPP-GC AMPAR-LTP, but heterosynaptic LPP-GC AMPAR-LTD. Heterosynaptic NMDAR-LTP required Ca^2+^ release from internal stores, implicating long-range signaling via the ER. Furthermore, homosynaptic and heterosynaptic NMDAR-LTP increased the sensitivity to GluN2D antagonism, suggesting the synaptic recruitment of GluN2D-containing NMDARs as expression mechanism. Collectively, our findings describe a novel activity-dependent synaptic mechanism whereby distinct entorhinal inputs interact functionally to regulate the dentate gyrus output.

### Activity induces homosynaptic NMDAR-LTP at MPP-GC but not at LPP-GC synapses

BTDP of NMDAR-mediated transmission was previously reported in midbrain dopamine neurons (Harnett et al., 2009) and in GC-CA3 synapses (Hunt et al., 2013). *In vivo* recordings showed that medial entorhinal cortex neurons fire bursts of action potentials (Latuske et al., 2015; Ebbesen et al., 2016; Csordas et al., 2020), and GCs which normally fire sparsely, can generate high frequency bursts of back propagating action potentials (Pernia-Andrade and Jonas, 2014; Diamantaki et al., 2016; Senzai and Buzsaki, 2017). In the present study, we found that pairing pre and postsynaptic burst activity selectively induced homosynaptic NMDAR-LTP at MPP-GC. Consistent with previous findings, this NMDAR-LTP was postsynaptically expressed, and required a rise in postsynaptic Ca^2+^ (Hunt and Castillo, 2012), and NMDAR and mGluR5 co-activation (Jia et al., 1998; Harney et al., 2006; Kwon and Castillo, 2008; Rebola et al., 2008; Harnett et al., 2009). In contrast, the BTDP paradigm did not induce homosynaptic plasticity at LPP-GC synapses. This finding might be explained by a strong dendritic attenuation of backpropagating action potentials as a function of distance from the soma (Krueppel et al., 2011; Kim et al., 2018). Also, as in distal synapses of other pyramidal neurons (Larkum et al., 2001; Sjostrom and Nelson, 2002), backpropagating action potentials may not be sufficient to unblock NMDARs at LPP-GCs. In a recent study, a more standard pre-post pairing protocol (i.e., single action potentials and synaptic responses) also failed to elicit LPP-GC AMPAR plasticity, whereas this plasticity was induced with a theta-burst stimulation of LPP inputs that generated local dendritic spikes (Kim et al., 2018). Our data focusing on NMDAR-mediated transmission support the idea that synaptic plasticity rules at proximal and distal inputs onto GCs differ.

### MPP-GC burst activity triggers opposite heterosynaptic LPP-GC plasticity of NMDAR- and AMPAR-mediated transmission

To our knowledge, we provide the first evidence for heterosynaptic NMDAR plasticity. Given the unique properties of the NMDAR, which can result in signal amplification, temporal summation, and increased Ca^2+^ influx (Hunt and Castillo, 2012), it is expected that NMDAR potentiation modifies the dendritic properties of GCs at distal LPP inputs during the expression of heterosynaptic NMDAR LTP. Unlike NMDAR-mediated transmission, AMPAR potentiation at MPP-GC synapses was accompanied by heterosynaptic AMPAR-LTD at LPP-GC synapses. Similar heterosynaptic LTD of AMPAR-mediated transmission has been reported before at many other synapses (Lynch et al., 1977; Chistiakova et al., 2014; Chater and Goda, 2021). Our findings provide further evidence that NMDARs and AMPARs have opposite heterosynaptic plasticity rules, which might impact dendritic integration by GCs.

### Homosynaptic and heterosynaptic NMDAR-LTP are mediated by GluN2D recruitment

A role for GluN2D-containing NMDARs in plasticity has been previously described at several synapses (Harney et al., 2008; Krause et al., 2008; Dubois and Liu, 2021; Eapen et al., 2021). As reported at MPP-GC synapses (Harney et al., 2008), we found that the maintenance of both homosynaptic MPP-GC and heterosynaptic LPP-GC NMDAR-LTP were blocked by GluN2D-subunit antagonists, suggesting that both forms of plasticity require recruitment of GluN2D-containing receptors to the synapse, presumably forming di- (GluN1/GluN2D) or tri- (GluN1/GluN2D/GluN2B) heteromers. In any case, the functional properties differ from those of diheteromeric GluN1/2B receptors (Yi et al., 2019). While the recruitment of heteromeric GluN2D-containing receptors could result in a NMDAR-EPSC with a slower decay (Cull-Candy et al., 2001; Hansen and Traynelis, 2011; Sanz-Clemente et al., 2013), no differences in NMDAR-EPSC decay were observed before and after LTP (Harney et al., 2008). Importantly, an important property of GluN2D-containing diheteromeric receptors is the low sensitivity to Mg^2+^ blockade relative to GluN2A or GluN2B-containing receptors (Paoletti et al., 2013). Although at distal synapses backpropagating APs may not be sufficient to remove the NMDAR Mg^2+^ block (Larkum et al., 2001; Sjostrom and Nelson, 2002), expression of GluN2D-containing NMDARs after NMDAR-LTP induction may be one factor that helps explain why NMDARs exhibit different synaptic plasticity learning rules than AMPARs. Notably, heterosynaptic changes in NMDAR subunit composition have been previously reported in CA1 neurons (Han and Heinemann, 2013). GluN2D-mediated synaptic transmission has been reported in hippocampal interneurons of adult mice (von Engelhardt et al., 2015), but whether activity modulates the expression of synaptic GluN2D-containing NMDARs is unclear. An increase in GluN2D subunit recruitment, which is expected to modify NMDAR properties and function (Yashiro and Philpot, 2008; Paoletti et al., 2013), suggests that long-term potentiation of NMDAR-mediated transmission could impact other forms of Hebbian plasticity.

### Homosynaptic vs. heterosynaptic NMDAR-LTP have unique Ca^2+^ requirements

Mechanistically, the induction and expression of NMDAR plasticity share common properties across synapses (Rebola et al., 2010; Hunt and Castillo, 2012), including postsynaptic NMDAR-mediated Ca^2+^ influx and Ca^2+^ release from internal stores. Our data shows that while homosynaptic NMDAR-LTP requires a rise in postsynaptic Ca^2+^, it is insensitive to pharmacological depletion of internal Ca^2+^ stores, suggesting that at MPP-GC synapses the postsynaptic Ca^2+^ rise via NMDARs and action potential firing is sufficient to induce plasticity. In contrast, we found that depletion of Ca^2+^ internal stores abolished heterosynaptic NMDAR-LTP, and that internal Ca^2+^ release from stores is mediated by ryanodine receptors. These data strongly suggest that homosynaptic NMDAR-LTP can change the synaptic rules at distal LPP-GC synapses, in part by eliciting the release of internal ER Ca^2+^ stores. While homosynaptic NMDAR-LTP at MF-CA3 synapses requires IP3-dependent Ca^2+^ release (Kwon and Castillo, 2008; Hunt et al., 2013), there is evidence in the amygdala that heterosynaptic, but not homosynaptic plasticity requires ryanodine-dependent Ca^2+^ release from internal stores (Royer and Pare, 2003), and ryanodine-dependent Ca^2+^ release can significantly alter the induction requirements of synaptic plasticity in GCs (Wang et al., 1996). Additionally, in mesolimbic dopamine neurons exposed to alcohol, increased susceptibility to induce NMDAR-LTP is dependent on Ca^2+^ release from ER stores (Bernier et al., 2011). Notably, EM studies of pyramidal hippocampal neurons show that the smooth ER extends to distal dendrites and dendritic spines (Spacek and Harris, 1997). The coincident increase in Ca^2+^ release from the ER, along with increased GluN2D recruitment to ease the Mg^2+^ block, might provide an ideal scenario for the induction and expression of heterosynaptic NMDAR LTP at distal synapses. Future *in vivo* Ca^2+^ imaging studies in combination with genetic strategies might help resolve exactly how these two events converge to mediate heterosynaptic plasticity and potentially contribute to dentate gyrus-depending learning.

In summary, BTDP of NMDAR-mediated transmission at MPP synapses can exert long-term, powerful control over NMDAR transmission at distal LPP-GC synapses. Dynamic changes in NMDAR transmission could shift the induction threshold of NMDAR-dependent forms of plasticity, as previously observed in CA3 neurons, where NMDARs can selectively adjust synapses in a heterosynaptic manner (Tsukamoto et al., 2003), act as a metaplastic switch for AMPAR-mediated plasticity (Astori et al., 2010; Rebola et al., 2011) and contribute to metaplasticity of AMPAR LTP (Hunt et al., 2013). Further work is required to determine whether homo- and heterosynaptic NMDAR potentiation occurs in behaving animals, and its contribution to dentate gyrus-dependent memory processes such as pattern separation (McHugh et al., 2007).

## Acknowledgments

This research was supported by the National Institutes of Health: R01-MH125772, R01-NS113600 and R01-MH081935 to PEC. ARR was partially supported by the Brain & Behavior Research Foundation Young Investigator Award (2014) and the Ford Foundation Postdoctoral Fellowship (2013). KN was partially supported by the American Epilepsy Society Postdoctoral Research Fellowship (2020). M.C was supported by NIH R25GM104547. We thank all Castillo Lab members for helpful discussions

